# A Framework for Relating Natural Movement to Length and Quality of Life in Human and Non-human Animals

**DOI:** 10.1101/2022.11.28.518240

**Authors:** Iain Hunter, Raz Leib

## Abstract

Natural movement is clearly related to health, however, it is also highly complex and difficult to measure. Most attempts to measure it focus on functional movements in humans, and while this a valid and popular approach, assays focussed on particular movements cannot capture the range of natural movement that occurs outside them. It is also difficult to use current techniques to compare movement across animal species. Interspecies comparison may be useful for identifying conserved biomechanical and/ or computational principles of movement that could inform human and veterinary medicine, plus several other fields of research. It is therefore important that research develops a system for quantifying movement in freely moving animals in natural environments and relating it to length and quality of life (LQOL). The present text proposes a novel theoretical framework for doing so, based on movement ability (*MA*). *MA* is comprised of three major variables – Movement Quality, Movement Complexity, and Movement Quantity – that may represent the most important components of movement as it relates to LQOL. A constrained version of the framework is validated in *Drosophila*, which suggests that *MA* may indeed represent a useful new paradigm for understanding the relationship between movement and length and quality of life.

## 1 Introduction

Movement is fundamental to life, and it can take many forms. This work is concerned with large-scale, skeletal-level movement, and its measurement to support length and quality of life (LQOL). This type of movement is inherent in changes in joint angle, such as shoulder flexion, which contributes to a wide range of important behaviours (Gill *et al*., 2020). This includes reaching, which supports feeding in many animals (Whishaw *et al*., 2017).

A body of research has investigated large-scale movement and how it may support length and/ or quality of life in humans (Király and Gondos, 2014; Heinrich *et al*., 2015; Cholewa *et al*., 2014). This work often constrains movement to simplify or enable understanding of it. For example, there have been many instances where assessment is particular to the sort of reaching described above (Wagner *et al*., 2007; Paolucci *et al*., 2021) or another, ‘functional’ movement in isolation of further movement complexity (Wagner *et al*., 2014; Slaughter *et al*., 2015; Dogu *et al*., 2013). Experiments that measure non-human animal movement usually take a similar, or more abstracted approach in cases where research questions are concerned with conservation (Katzner and Arlettaz, 2020). Such work is important, however, it cannot capture much of what most natural movement represents: a highly complex, dynamic, and often whole body output that is a probabilistic function of an animal’s physiology and environment (Goossens *et al*., 2020; Latash and Anson, 1996). The present text describes a novel theoretical framework within which it may be possible to capture some of the complexity and dynamism of natural movement, and relates it to LQOL, in a way that is not possible using traditional protocols that dissect it.

The framework presented below has the potential to form the basis of a model. In addition to quantifying movement and relating it to LQOL, it represents a single system that may be used to compare natural movement both within and between species of animal (intraspecies, e.g., human to human, or interspecies, e.g., human to canid, respectively). This type of comparison is important; it may enable the identification of general (unifying) biomechanical and/ or computational principles of movement conserved across animals, which could form the principal components of locomotion that should be addressed first when treating movement disorder. These principles may also be used to realise more animal-like movement in robots (Kashiri *et al*., 2018), or to describe how sensory information is processed to provide appropriate action selection (reviewed in (Franklin and Wolpert, 2011)) which could inform control theory (Zarandi and Mosadegh, 2016). Finally, interspecies comparison of natural movement may also support veterinary and human medicine, conservation (Wright *et al*., 2020), and improve understanding of predator-prey dynamics (Keim *et al*., 2021).

The framework is written to express a human or non-human animal’s natural movement and the extent to which it supports LQOL, as Movement Ability (*MA*). *MA* is defined as a function of three variables: Movement Quality (*M*_*qual*_), Movement Complexity (*M*_*comp*_) and Movement Quantity (*M*_*quant*_). Each relates to different aspects of natural movement and is derived from several more, which are described conceptually and mathematically, in the Framework section. They are also explored in greater depth in the Discussion (section 4).

## 2 Framework

### 2.1 Movement Ability

It is proposed that a human or non-human animal’s movement ability is measured as Movement Ability (*MA*):

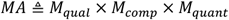

Where:

- *M*_*qual*_= Movement Quality
- *M*_*comp*_ = Movement Complexity
- *M*_*quant*_ = Movement Quantity

Exemplar calculations of these variables are given at the end of each of the relevant sections below. Note that while these examples are based on humans (*MA*_*human*_ (healthy) or *MA*_*humanMD*_ (movement disorder)) or mice (*MA*_*mouse*_ (healthy) or *MA*_*mouseMD*_ (movement disorder)), the calculations are the same regardless of species, and may be applied to any human or non-human animal. Indeed, we validate a constrained version of the framework in the fruit fly, *Drosophila melanogaster*, below (Results, section 3).

### 2.2 Movement Quality

Movement Quality (*M*_*qual*_) is designed to score a human or non-human animal’s movement to reflect the absence or presence, and probability of future development of movement dysfunction. High *M*_*qual*_ therefore suggests absence and low probability of future dysfunction, while low *M*_*qual*_ represents the opposite (functional disadvantage). It is expressed as:

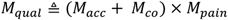

Where:

*M*_*acc*_ = movement accuracy scored from 0-1. Based on decimal of percentage of recording time (*rt*), in which joint angle(s) of all major joints exist within their safe ranges (1 = 100%, 0.5 = 50% etc.), while changing angle to an extent and with a rate and directional variability consistent with normal motor control.

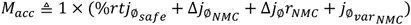

Joint (*j*) is defined as the centre of the fulcrum responsible for articulation of a body part, which contributes to movement or locomotion of human or non-human animal. Joint angle (*j*_∅_) is measured as three-dimensional angle (∅) formed between positions of *j* and most distal component of body part in x, y and z axes of space (*x, y and z* below), where it either forms another joint (e.g. hip to knee at femur) or terminates to complete a particular part of the body plan (e.g. atlanto-occipital joint to nose):

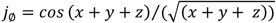

Safe joint angle 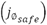 is defined as *j*_∅_ which is equal to or less that the anatomically determined range appropriate to a given joint (*j*_*anat*_), e.g., *x*_0°_ − *x*_180°_ for a hinge joint:

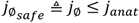

Extent of change in joint angle is measured as the difference in three-dimensional joint angle (Δ*j*_∅_, a vector) of a given joint, which occurs between two points in recording time:

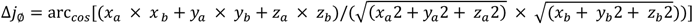

Extent of change in joint angle consistent with normal control of movement 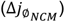 is defined as that which occurs within ± 2 standard deviations (2σ) of the mean unidirectional displacement of that joint in a sample of healthy human or non-human animals of the same species. It is therefore defined as:

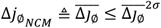

Rate of change in joint angle is measured as the velocity with which *j*_∅_ changes (Δ*j*_∅_*r*, t = time).

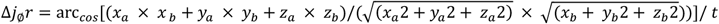

Rate of change in joint angle consistent with normal control of movement (Δ*j*_∅_*r*_*NCM*_) is defined as that which occurs within ± 2 standard deviations (2σ) of the mean rate of change recorded in a sample of healthy human or non-human animals of the same species:

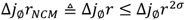

Directional variability is expressed as variance of joint angle 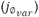 observed during recording time. Thus, directional variability consistent with normal control of movement 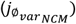, is the variance of *j*_∅_ that is less than or equal to the variance recorded in a sample of healthy human or non-human animals of the same species (σ^2^):

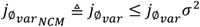

*M*_*co*_= movement coordination is scored from 0-1, based on decimal expression of percentage of recording time (*rt*) in which the relative positions of two or more joints are consistent with normally coordinated movement (1 = 100%, 0.5 = 50% etc.)

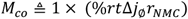

The relative positions of two joints (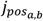, where *a* and *b* define different joints) is expressed in a similar way to Δ*j*_∅_. It is:

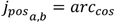

When relating the positions of more than two joints 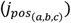, the shared angle is expressed using a scalar product form formula to find the Cartesian equation of the plane formed by those joints:

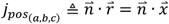

Where 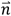 is any vector normal to the plane and 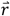 is a generic vector in 3-dimensional space (e.g. *q, r, c*). Relative position of two or more joints consistent with normally coordinated movement 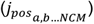 describes that which occurs within ± 2 standard deviations (2σσ) of the mean rate of change recorded in a sample of healthy human or non-human animals of the same species:

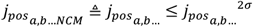

*M*_*pain*_= likelihood of pain present in movement, based on avoidance scored from 0-1. Specifically, it is quantified as the inverse decimal of the percentage of avoidance of a behaviour correlated with a human or non-human animal’s survival *M*_*pain*(*x*)_, versus its mean occurrence in a healthy sample:

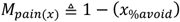

Behaviour(s) correlated with a human or non-human animal’s survival are defined by its species-specific ethogram. All these behaviours are scored for *M*_*pain*_, however, only the lowest scoring (i.e. largest avoidance, given as *x* below) is entered into the equation for *M*_*qual*_. *M*_*pain*_may, therefore, be expressed as:

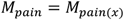

In summary, assuming a recording time of 24h and arbitrary units, *M*_*qual*_may be scored as:

- (0.9 + 0.9) × 1 = 1.8, for 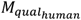 that moves with a high degree of accuracy and coordination, without avoiding behaviour correlated with survival.
- (0.51 + 0.6) × 0.8 = 0.96, for 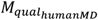 that moves with a relatively low degree of accuracy and coordination, while occasionally avoiding behaviour correlated with survival.
- (0.96 + 0.94) × 1 = 1.9, for 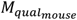 that moves with a slightly greater degree of accuracy and coordination than the *MA*_*human*_, without avoiding behaviour correlated with survival.
- (0.48 + 0.66) × 0.7 = 0.86, for 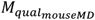 that moves with a low degree of accuracy and coordination that is similar to human movement disorder, while avoiding behaviour correlated with survival more regularly.

Finally, *M*_*qual*_ is normalised to the highest value produced by an individual in a sample, to give a modified z-score that enables expressing *M*_*qual*_ as a value from 0-1.

### 2.3 Movement Complexity

Movement Complexity (*M*_*comp*_) expresses the complexity inherent in human or non-human animal’s movement. High *M*_*comp*_ represents movement that is varied and difficult to predict, given the series of (other) movements performed prior to it. Low *M*_*comp*_ reflects more repetitive and predictable movement. It is defined as:

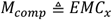

Where:

*EME*= Effective Measure Complexity (Grassberger, 1986), and *x* is a sequence of alignments. Alignment (*a*) is the momentary position of a human or non-human animal during movement, measured as the orientation of its rostrocaudal axis in three-dimensional space, plus the orientation of head and limbs (where they are present) relative to the body. *EMC*_*x*_, therefore, calculates the ability to accurately predict the occurrence of a particular alignment (*a* within *x*, i.e., according to the series of (other) alignments that precede it. It may be expressed as follows:

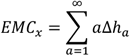

Where *h*_*a*_ is a ‘new’ alignment. Note that alignments are converted into symbols to be processed as a string, as is common in the field.

Thus, assuming a total recording time of 24h and arbitrary units, *M*_*comp*_ may be scored as:

- 100 for 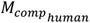 that conducts a weekly tumbling practice, which produces a complexity score that is considerably above the human population mean.
- 38 for 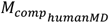 that changes alignment only as much as is necessary to perform basic movements to support survival.
- 135 for 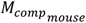 that produces highly complex movement (e.g. climbing, inverting and compression) as part of navigating an environment which demands it.

23 for 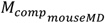 that cannot change alignment as its heathy counterpart can, so, like 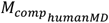, only does so as much as is necessary to support survival.

Like *M*_*qual*_, *M*_*comp*_ is normalised to the highest value produced by an individual in a sample, to enable expressing *M*_*qual*_as a value from 0-1.

### 2.4 Movement Quantity

Movement Quantity (*M*_*quant*_) reflects the amount of movement a human or non-human animal produces during recording time. Specifically, it considers the range of movement velocity and total displacement observed. Thus, high *M*_*quant*_ may indicate that a human or non-human animal has the capacity to move with a high maximum velocity, over long distances. It is expressed as follows:

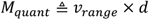

Where:

*υ*_*range*_ is range of movement velocity, calculated from the highest and lowest non-idle centre of mass velocities observed during recording time:

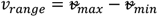

*d* is the total distance travelled by the animal’s centre of mass (*c. c.m*._*x*_− *c. c.m*._*y*_) as defined by the limits of the sum (*c. c.m*. = 1→*c. c.m*.∞):

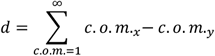

Assuming recording time of 24h, *M*_*quant*_ may be scored (using m/ s for *υυ*_*range*_ and km for *aa*):

- 3.35 × 12 = 40.2, for *MA*_*human*_ that ran 10km, and then was idle for most of the day.
- 2 × 4 = 8, for *MA*_*humanMD*_ that only moved a relatively short distance, slowly.
- 2 × 6 = 12, for *MA*_*mouse*_ that was highly active, but that did not move with as much velocity or as far as the *MA*_*human*_.
- 0.17 × 2 = 0.34, for *MA*_*mouseMD*_ that like the *MA*_*humanMD*_, only moved a very short distance, slowly.

As for *M*_*qual*_and *M*_*comp*_, *M*_*quant*_ is normalised to the highest value produced by an individual in a sample, so that is expressed as a value from 0-1.

### 2.5 Calculating Movement Ability

Combining the exemplar calculations of *M*_*qual*_, *M*_*comp*_ and *M*_*quant*_, and assuming normalisation to arbitrary values that represent the highest score in sample, *MA* may be given as follows:

- *MA*_*human*_ = 0.8 × 0.7 × 0.8 = 0.49
- *MA*_*humanMD* =_ 0.4 × 0.3 × 0.5 = 0.06
- *MA*_*mouse* =_ 0.7 × 0.8 × 0.6 = 0.34
- *MA*_*mouseMD* =_ 0.3 × 0.3 × 0.2 = 0.02

## 3 Results

While the exemplar calculations given above provide important insight into the way *MA* is constructed, it was important to test the framework with data generated in experiments. Consequently, a constrained version of the framework was tested on adult male *Drosophila* (4d after eclosion) of three genotypes, which we hypothesised may demonstrate different *MA*. These genotypes were *Canton-S* (*CS*, wild type control), *bang-sensitive* (*bss*) and *inactive*^*3621*^ (*iav*). *Bss* animals express a mutation that leads to hyperactivity in the *Drosophila* voltage-gated sodium channel (Parker *et al*., 2011) and are commonly used as a model of epilepsy (Dare *et al*., 2021; Parker *et al*., 2011). *iav* mutants lose function in an ion channel that contributes to mechanotransduction in proprioceptors (Gong *et al*., 2004; Albert and Gopfert, 2015; Fushiki, Kohsaka and Nose, 2013), so may demonstrate dysfunctional movement (Fushiki, Kohsaka and Nose, 2013).

*M*_*comp*_, *M*_*quant*_, and *M*_*comp*_ × *M*_*quant*_ (a version of *MA* constrained by omission of *M*_*qual*_) were calculated for each genotype from video recordings of animals moving freely inside an arena (Figure 1A-B), and each of these parameters was related to the lifespan of life of the different genotypes (Figure 1D-F). *M*_*qual*_ and quality of life could not be calculated due to limitations addressed in the Discussion.

**Figure 1:**
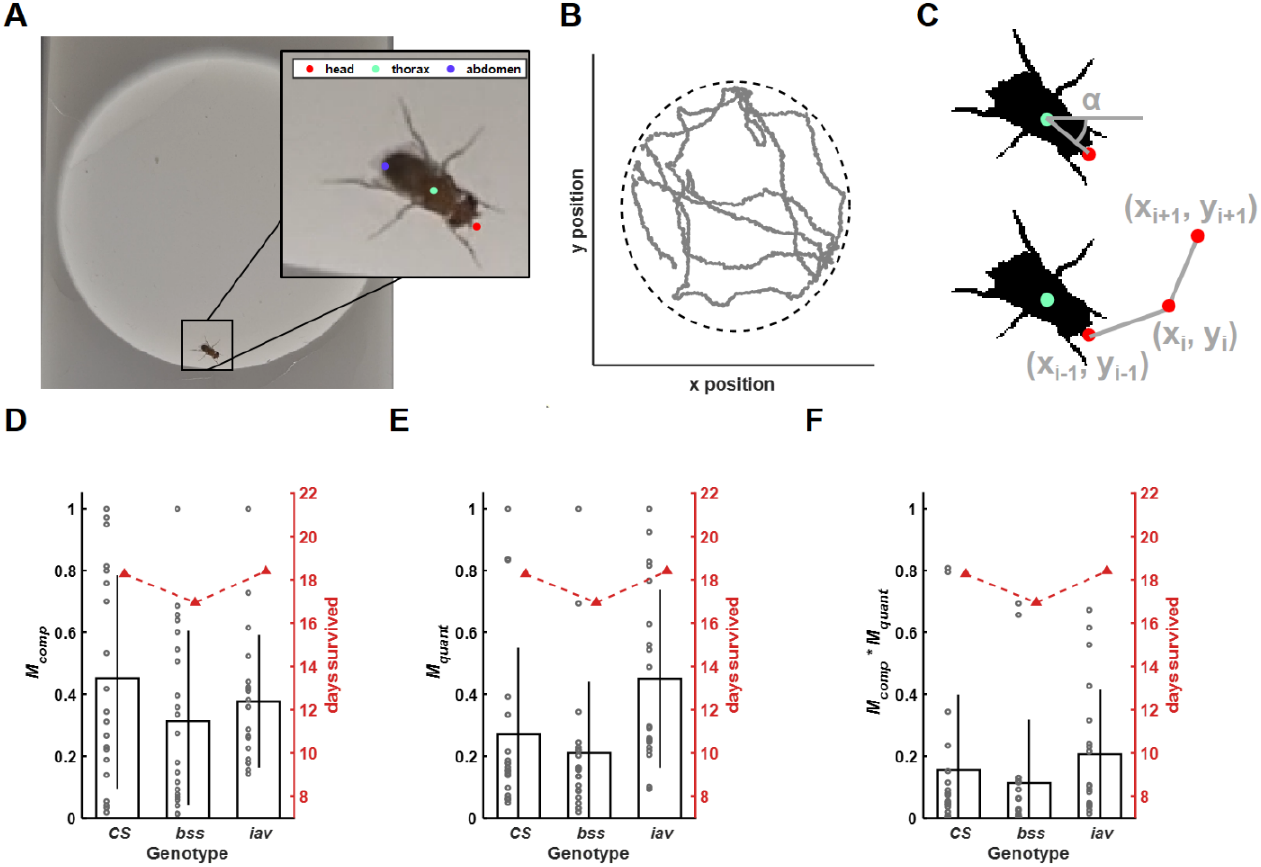
Analysis of Movement Ability in Drosophila. *(A) experimental setup: adult male Drosophila moved freely inside an arena. This movement was recorded on camera, and analysed to produce traces that followed the head, thorax and abdomen, using DeepLabCut (Mathis et al*., *2018). (B) representative trace generated by tracking the thorax of a fly (head and abdomen were also recorded, but are not shown for simplicity). Dashed black line indicates the edge of the arena. (C) Movement Complexity* (*M*_*comp*_) *was calculated using alignments based on the angle of the vector connecting the head and thorax, with respect to the horizon line (upper panel). Movement Quantity (M*_*quant*_) *was calculated as the product of the range of velocity and total distance traveled during recording time (bottom panel). Both were normalized to the highest score recorded for that variable (e.g. all M*_*comp*_ *values were normalized to highest M*_*comp*_*) so that they could be scored from 0-1. (D-F) graphs* relating *M*_*quant*_, and *M*_*comp*_ × *M*_*quant*_, to the mean number of days that flies from 3 genotypes (Canton-S, CS; bang sensitive, bss; inactive, iav) survived. Black bars show means, circles individual values, black lines are standard deviations and red triangles are mean days survived after experiments.

*M*_*comp*_ was highest in *CS* (0.45 ± 0.35, *n* = 19), and reduced in the two mutants (*bss*: 0.31 ± 0.29, *n* = 19; *iav*: 0.37 ± 0.21, *n* = 20). Differences in *M*_*comp*_ were not significant (F2,34.61 = 0.864, *p* = 0.43, by 3-way ANOVA with multiple comparisons), however, differences in *M*_*comp*_ appeared similar to differences in lifespan. Specifically, *CS* and *iav* moved with more *M*_*comp*_ and had a longer lifespan (*CS*: 18.2 ± 13d, *n* = 19; *iav*: 18.4 ± 9.8d, *n* = 20) than *bss* (16.9 ± 9d, *n* = 19). Testing the relationship between *M*_*comp*_ and lifespan within the three gentypes, revealed a strong correlation in *CS* (R^2^ = 0.61, *p* < 0.001). There was no clear correlation in either mutant genotype (*bss*: R^2^ = 0.19, *p* = 0.06; *iav*: R^2^ = 0.004, *p* = 0.79), which likely reflects the large variability observed in the behaviour of the mutant flies. Indeed, it is well established that mutant animals behave with more variability than wild types. Thus, the results we generated in *CS* validate using *M*_*comp*_ to relate movement to LQOL. Those generated in mutants propose that a much larger sample size should be used to enable greater power for validating *M*_*comp*_ in these and similar genotypes in future.

Results for *M*_*quant*_ were reminiscent of those for *M*_*comp*_, however, there were some important differences. *M*_*quant*_ was higher for *CS* and *iav* (*CS*: 0.27 ± 0.29, *n* = 19; *iav*: 0.44 ± 0.28, *n* = 20) than *bss* (0.21 ± 0.24, *n* = 19). There was a significant difference in *M*_*quant*_ (F2,57 = 4.01, *p* =0.024), post-hoc analysis showed that this was due to a significant difference between *bss* and *iav* (*bss* vs. *iav*: t55 = 2.7, *p* = 0.027) but not between *CS* and *bss* or *iav* (*CS* vs. *bss*: t55 = 0.67, *p* = 1.0; *CS* vs. *iav*: t55 = 2.03, *p* = 0.14). These values trended in the same direction as lifespan and yet in contrast to *M*_*comp*_, there was no clear evidence of a correlation between *M*_*quant*_ and length of life for any of the genotypes (*CS*: R^2^ = 0.11, *p* = 0.15; *bss*: R^2^ = 0.003, *p* = 0.8; *iav*: R^2^ < 0.01, *p* = 0.92). This was an unexpected result for *CS* (less for *bss* and *iav* due to the variability in behaviour associated with mutation), given the evidence that suggests moving more extends life (Reimers, Knapp and K., 2012; Lee *et al*., 2022). It is likely that recordings (180s) were too short to capture sufficient data for *M*_*quant*_ to relate it to LQOL, and so poses that longer recordings should be used to validate this aspect of the framework.

As expected, *M*_*comp*_ × *M*_*quant*_ (i.e., *MA* excluding *M*_*qual*_) followed similar trends to those observed for individual variables. *M*_*comp*_ × *M*_*quant*_ was higher for *CS* and *iav* (*CS*: 0.15 ± 0.24, *n =* 19; *iav*: 0.2 ± 0.2, *n =* 20), and lower for *bss* (0.11 ± 0.2, *n =* 19). *M*_*comp*_ × *M*_*quant*_ was correlated with lifespan in *CS* (R^2^ = 0.37, *p* = 0.005), which validates using *MA* to relate lifespan and LQOL. While this relationship was not apparent in either mutant genotype (*bss*: R^2^ = 0.002, *p* = 0.84; *iav*: R^2^ = 0.002, *p* =0.83), the trends we observed between the primary variables (*M*_*comp*_ and *M*_*quant*_) or *M*_*comp*_ × *M*_*quant*_ and lifespan, plus the convincing correlations observed for *CS*, suggest that it exists in the mutants too. As mentioned previously, future work could test this hypothesis in experiments with greater power (i.e., a larger sample size).

## 4 Discussion

The hypothetical (exemplar) scores for *MA* and each of its major variables (*M*_*qual*_, *M*_*comp*_ and *M*_*quant*_) indicate the potential utility of the framework. Specifically, the higher score for the healthy human (0.49) and lower for the movement disorder patient (0.06) proposes that *MA* may be used to relate natural movement to LQOL. The examples also demonstrate that comparisons may be drawn between different animals of the same species (0.34 versus 0.02 for healthy and movement disorder mice, respectively), and between different species (0.49 versus 0.34 for a healthy human and mouse, respectively). The results generated while testing the model support these examples. Specifically, the correlations between *M*_*comp*_ and *M*_*comp*_ × *M*_*quant*_ and lifespan in *CS*, propose that *MA* is a valid means of relating natural movement to LQOL. The trends towards the same relationships in *bss* and *iav* suggest that further support could be provided by demonstrating significant correlations are significant in experiments using larger sample sizes. This is noteworthy because as mentioned previously, intraspecies and interspecies comparisons may be useful in identifying unifying principles of movement. They could inform fields concerned with medicine and veterinary medicine/ rehabilitation (Tarakad, 2020), optimising training programs designed to increase LQOL in already healthy human and non-human animals, robotics (Aguilar *et al*., 2016), action selection, control theory (Zarandi and Mosadegh, 2016), conservation (Wright et al., 2020) and understanding predator/ prey dynamics (Keim et al., 2021).

Future work should continue to develop the potential to use *MA* to study the relationship between movement and LQOL, through further validation of the model. It should test the hypothesis that *M*_*qual*_and by extension, *M*_*qual*_× *M*_*comp*_ × *M*_*quant*_, correlates with LQOL. It was not possible to test *M*_*qual*_in the experiments described in the present text because most flies spent some time walking along the side of the behavioural arena. This obscured their legs from the view of the camera and made it difficult to perform the calculations necessary to compute *M*_*qual*_. This issue could be addressed using a tool such as Anipose (predicts 3D pose based on 2D information (Karashchuk *et al*., 2021)) or by using several cameras in a larger arena, so that leg position is reported for the duration of the recording.

Alternatively, or in addition to recording behaviour in *Drosophila*, further validation could test the ability to measure *MA* in other species. For example, movement of freely-moving wild or semi-wild animals, such as those being monitored by the St Kilda Soay Sheep, Isle of Rum Red Deer or Isle Royale Wolf projects, could be recorded by drone or wearable sensors, and analysed according to the framework. Such projects usually record the length of life of the animals they monitor, so *MA* could be related to LQOL. *MA* could also be related to LQOL in human participants who have had their movements recorded on video, by a wearable sensor, or in a living lab. While early phases of validation may use experiments subject to certain limitations (e.g., the relatively short recordings associated with monitoring human movement in a living lab), testing should continue until *MA* has been validated by recording movement of human or non-human animals in natural environments for extended periods of time (e.g. 24h-1w). This duration is important, as longer recordings are more likely than shorter, to capture the complete spectrum of movements an animal performs. However, validating *MA* over extended timeframes would take considerable effort. It would require ensuring reasonably unobstructed, continuous monitoring of behaviour for the duration of a trial, and the large amounts of data generated would have to be stored and analysed. However, the potential utility of the framework, which is illustrated by the correlations described in the present text, suggest this effort may be worthwhile.

It is important to highlight that the potential utility of the framework is enabled by its novel structure. As stated in the Introduction, it is difficult to measure ‘complete’ natural movement. Traditional approaches are, therefore, highly constrained by design. The *MA* model affords significantly greater flexibility because it is based on “higher order” principles that reflect commonalities of movement, which are not limited by peculiarities of individual animals or even species. These principles are defined in the primary variables of *M*_*qual*_, *M*_*comp*_ and *M*_*quant*_, and it is possible that they are broad enough for all parameters that describe natural movement to fall under them. This important idea – that *MA* (as quality, complexity and quantity of movement) may be a unifying principle sufficient to explain differences in movement between animals – must be tested in future work. This work should address the question: are there any variables or components of movement which cannot reasonably fit under the headings of Movement Quality, Movement Complexity and Movement Quantity? One such variable could be movement relevance. That is, the degree to which movement supports survival or reproductive fitness. While the current version of the model accounts for avoidance of behaviours related to survival as evidence for pain in movement, there may be utility in weighting *M*_*qual*_to favour behaviour correlated with survival or reproductive fitness. Notably, scoring both avoidance or performance of behaviour correlated with survival, depends on establishing those behaviours in ethograms. Robust ethograms have been described for certain animals (e.g. mouse (Garner, 2023)), however, establishing them is difficult. Machine learning tools such as DeepEthogram (Bohnslav *et al*., 2021) may help expedite creating ethograms, so that they are very beneficial to calculating *MA*.

Finally, given their novelty and importance to MA, it is necessary to discuss how the authors decided to include and define the three primary variables (*M*_*qual*_, *M*_*comp*_ and *M*_*quant*_). *M*_*qual*_was selected to reflect research that suggests abnormal biomechanics and pain during movement, is associated with lower quality (and perhaps length) of life. This includes work which shows that disordered movement is related to injury (Lohman *et al*., 2019) and is symptomatic of movement disorder (Morris *et al*., 2001). Similarly, physiotherapy research aims to improve an established practice that, in essence, aims to treat disordered (i.e. low quality) movement to improve quality of life (Alphonsus, Su and D’Arcy, 2019). Most prior attempts to quantify movement quality, which (arguably) include the functional movement screen (FMS), are: (1) constrained to human functional movements; (2) subject to notable controversy regarding their effectiveness (Frost *et al*., 2012; Beardsley and Contreras, 2014). The parameters chosen to define *M*_*qual*_here were, therefore, based on what seemed to be sensible relationships relating the biomechanical accuracy, coordination and pain experienced during movement to LQOL (Lu and Chang, 2012; Dunsky, 2019; Ferrell, 1995). It is important to note that despite them being logical, the nature of these relationships are open to debate. This is particularly true in the case of relating pain to movement (i.e. *M*_*pain*_), as it is predicated on accurate measurement of an experience that is at least somewhat subjective (Breivik *et al*., 2008). Indeed, many popular means of scoring pain (such as using grimace scores (Cohen and Beths, 2020)) cannot be used for calculating *M*_*pain*_, as *MA* is designed to enable interspecies comparisons that are not supported in certain animals that do not possess the features necessary to grimace. In attempting to measure pain while solving this problem, however, the framework may contribute something useful to the field.

Specifically, *M*_*pain*_ expresses pain as a percentage of avoidance of behaviours correlated with survival, with the latter identified in an animal’s ethogram. This method, which to the author’s knowledge is novel, could provide the basis for an objective score.

Movement Complexity (*M*_*comp*_) is clearly necessary for the survival of some animals (e.g. birds catching insects, or land animals evading predators), so it must feature in a system designed to relate movement to LQOL across species. Though the role that movement complexity plays in the lives of modern humans is less obvious, it seems likely that it contributes to LQOL. The ability to produce varied movement surely provides a person with opportunity that is unavailable to patients with disorders characterised by repetitive movement (Sanger *et al*., 2010).. It is therefore important to note that as for all the other components of movement discussed, it is very difficult to measure movement complexity. Indeed, much research is dedicated to how to define and quantify complexity generally (i.e., as a concept independent of its role in movement). It was some of this work that inspired defining *M*_*comp*_ using effective measure complexity (EMC (Grassberger, 1986)). In the context of the present text, EMC provides the ability to code alignments as symbols, and then to express the complexity of movement as the ability to predict a particular alignment occurring within a sequence of alignments. This novel application of an established paradigm, therefore, provides utility in this framework and could be used in other contexts. It may, for example, be valuable to compare the complexity of movement produced by Parkinson’s Disease patients following different treatments, to help decide which was most effective. Finally, it is difficult to design robots to move with the level of complexity (e.g., dexterity) present in animals (Xia *et al*., 2022), so *M*_*comp*_ may be used to test and improve it.

Movement Quantity (*M*_*quant*_) was included in the *MA* model due to its ability to reflect exercise volume, which, in turn, is associated with a longer lifespan (Reimers, Knapp and K., 2012; Lee *et al*., 2022). Exercise volume is also related to physical fitness, and arguably the best definition of this type of fitness is that of an animal or non-human animal’s ability to produce power (*work*/*time* (Knudson, 2009)). Reasonable extrapolation allows research to link power with LQOL (Lee, Hsieh and Paffenbarger, 1995), so that there is further precedent for measurements related to *M*_*quant*_ improving length and quality of life. It is important that *M*_*quant*_ is used in the framework instead of power, however, because calculating power artificially biases larger animals; they must work harder (produce more power) to move themselves the same distance as animals with less mass. This may affect the relationship of power with LQOL, as high mass due to poor body composition reduces LQOL in most animals (Toss *et al*., 2012). Thus, using displacement and range of velocity in the model instead, could provoke thought in researchers concerned with describing physical fitness. It may, for instance, inspire use of displacement and range of velocity, or a mass-adjusted version of power, in future assessments of fitness.

## 5 Conclusions

This work presents a model for calculating Movement Ability (*MA*), which is designed to measure movement of human and non-human animals in their natural environments, and relate it to length and quality of life. *MA* is comprised of three primary variables: *M*_*qual*_, *M*_*comp*_ and *M*_*quant*_, which may be sufficient for describing natural movement. Results generated by recording freely moving adult *Drosophila*, validate the proposed relationship between the model and LQOL, by demonstrating strong correlations between *M*_*comp*_ and *M*_*comp*_ × *M*_*quant*_ and lifespan. Other results suggest that *M*_*quant*_ could be similarly correlated with LQOL, and that this relationship would likely be demonstrable if certain limitations of the protocol employed in the present text are overcome. This work, therefore, introduces and validates a novel framework that can capture aspects of natural movement that more traditional means for measuring it cannot, while also relating it to length and quality of life. It could form the basis for further exploration and refinement of a paradigm with potential to make a meaningful contribution to the field.

## 6 Materials & Methods

### 6.1 Drosophila Rearing & Stocks

*Drosophila* stocks were kept on standard corn meal medium, at 25°C. Wild-type *Canton-S* or bang-sensitive (*para*^*bss1*^) were from lab stocks, *iav[3621]/C(1)DX, y[1] f[1]* was ordered from Bloomington Drosophila Stock Centre (BDSC, #52273).

### 6.2 Tracking Fly Movement

Male flies of different genotypes (*CS, bss* or *iav*) and similar age (∼4 after eclosion) were filmed (Samsung Galaxy S21 5G) moving freely in the chamber of a custom-made arena (Figure 1A), for 180s. Videos were uploaded to a computer, and movement data was extracted by a deep learningbased neural network model in DeepLabCut (Mathis *et al*., 2018). The model was trained by manual identification of three points on the body of each fly (the head, thorax, and abdomen (Figure 1A)) in 152 frames, from 8 randomly selected videos (19 frames per video). Once training was complete, the model was run so that it detected the three points in the rest of the frames, in all videos, automatically. The resulting output provided the pixel coordinates of each of the three markers (and so position of the fly in space (Figure 1B)), for the duration of the video. These coordinates were used to calculate *M*_*comp*_ and *M*_*quant*_.

*M*_*comp*_ was calculated by defining body alignment (*aa*) as the direction of the vector connecting the head and thorax of a fly (Figure 1C, top panel) in a single frame. The sum of changes in *aa* between all frames in each recording, was calculated using the *M*_*comp*_ formula given in the Framework section:

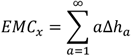

Once *M*_*comp*_ was calculated for each animal, values were normalised to that of the individual with the highest *M*_*comp*_, to enable expressing it from 0-1.

For *M*_*quant*_, range of velocity (*υ*_*range*_) was determined by subtracting the minimum (i.e. velocity > 0) from the maximum velocity observed during a recording. Velocity was calculated by subtracting the position of one point (thorax) in consecutive frames and dividing the result by 1/30, to account for the frame rate of the camera. Displacement (*aa*) was calculated as the total distance a fly moved during a recording, using the Euclidian distance one point (thorax) moved between consecutive frames (Figure 1C, bottom panel), to produce a distance in pixels. This distance was converted into Cartesian coordinates (mm) by multiplying it by a factor (5 × 10^4^), which accounted for the ratio of the camera resolution and distance from the arena/ fly. *υυ*_*range*_ and *aa* were multiplied, so that *M*_*quant*_was calculated as described in the formula given in the Framework section:

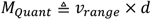

As for *M*_*comp*_, *M*_*quant*_ was calculated for each animal, then values were normalised to that of the individual with the highest score, to enable expressing it from 0-1.

### 6.3 Fly Survival

Flies used in tracking experiments were kept and placed in individual vials of standard corn meal medium, at 25°C. Flies were checked daily (excluding weekends) at ∼16:00, with deaths recorded in a spreadsheet (Microsoft Excel). Survivability was reported as days survived (specifically, days survived after movement tracking was complete), as opposed to according a more traditional measure such as after eclosion, for clarity.

### 6.4 Statistics

Statistical significance of differences in *M*_*quant*_, *M*_*comp*_ or *M*_*comp*_ × *M*_*quant*_ (dependent variables) between genotypes (independent variables) was assessed by 3-way ANOVA, with the threshold for significance set to *p* = 0.05. Homogeneity of variance was tested using the Levene statistic, and when homogeneity was violated, Welch’s F was used to calculate the F-ratio. Where differences were significant, post hoc analysis was performed using Bonferroni correction. All statistical analysis was performed in SPSS 20.

## 7 Statements and Declarations

Work on this project was supported by funding from the Wellcome Trust, awarded to Prof Richard Baines (217099/Z/19/Z). Declarations of interest: none.

## 8 Author Contributions

Iain Hunter: conceptualization, methodology, investigation, writing – original draft, project administration. Raz Lieb: formal analysis, visualization.

## 9 Acknowledgements

The authors thank Prof Richard Baines and Dr Cengiz Gunay, who provided valuable feedback on the text. They also thank Bramwell Coulson, for help conducting the survivability analysis.

